# Response-related signals increase confidence but not metacognitive performance

**DOI:** 10.1101/735712

**Authors:** Elisa Filevich, Christina Koß, Nathan Faivre

## Abstract

Confidence judgements are a central tool for research in metacognition. In a typical task, participants first perform perceptual (first-order) decisions and then rate their confidence in these decisions. The relationship between confidence and first-order accuracy is taken as measure of metacognitive performance. Confidence is often assumed to stem from decision-monitoring processes alone, but processes that co-occur with the first-order decision may also play a role in confidence formation. In fact, across a broad range of tasks, trials with quick reaction times to the first-order task are often judged with relatively higher confidence than those with slow responses. This robust finding suggests that confidence could be informed by a readout of reaction times in addition to decision-monitoring processes. To test this possibility, we assessed the contribution of response-related signals to confidence and, in particular, to metacognitive performance (i.e., a measure of the adequacy of these confidence judgements). In a factorial design, we measured the effect of making an overt (vs. covert) decision, as well as the effect of pairing a motor action to the stimulus about which the first-order decision is made. Against our expectations, we found no differences in overall confidence or metacognitive performance when first-order responses were covert as opposed to overt. Further, actions paired to visual stimuli presented led to higher confidence ratings, but did not affect metacognitive performance. These results suggest that some of the relationships between first-order decisional signals and confidence might indeed be correlational, and attributable to an upstream cognitive process, common to the two of them.

## Introduction

Confidence judgements about one’s own perception have been exploited in recent years as a particularly useful way to probe introspection (Fleming & Dolan, 2012). In what has become a standard paradigm, participants first make a perceptual judgement (first-order task) and immediately afterwards give a measure of confidence in their response (second-order task). The relationship between response accuracy in the first-order task and confidence in the second-order task is taken as a measure of *metacognitive performance*. A participant is said to have high metacognitive performance if she is able to assign high confidence exclusively to correct trials, but never, or seldom, to incorrect trials (Fleming & Lau, 2014). This standard paradigm has been used over a variety of domains of cognition from vision (Song et al., 2011) and audition (Ais, Zylberberg, Barttfeld, & Sigman, 2015), to memory (Baird, Smallwood, Gorgolewski, & Margulies, 2013) and value-based choice (De Martino, Fleming, Garrett, & Dolan, 2013). But it is still unclear what confidence reports actually represent, as the variables participants compute to generate them remain latent.

Under a normative view, confidence reports can be seen as a finer-grained description of the same perceptual evidence that leads to the binary first-order decision, and, specifically, correspond to the probability of giving a correct answer given the available perceptual discriminability (Pouget, Drugowitsch, & Kepecs, 2016; Sanders, Hangya, & Kepecs, 2016). In other words, whereas by experimental design participants are typically forced to choose between two options in the first-order task, they have the chance to more precisely describe the difficulty of their perceptual experience through confidence reports in the second-order task. In this view, introspection is required to produce accurate confidence reports. But recent results from correlational, modelling and brain stimulation approaches have challenged this standard view of confidence as description of perceptual evidence by showing that, beyond perceptual evidence, sensorimotor signals associated with the response provided to the first-order task may also contribute to confidence. At its simplest, this effect is manifest as a correlation over trials between first-order reaction times and confidence reports (e.g., (Charles, Chardin, & Haggard, 2018; Fleming, Weil, Nagy, Dolan, & Rees, 2010; Patel, Fleming, & Kilner, 2012). Further, it was shown that metacognitive performance was better in participants with large differences in response times between correct and incorrect responses (Faivre, Filevich, Solovey, Kühn, & Blanke, 2017). Beyond behaviour alone, correlational analyses showed that confidence is higher in the presence of sub-threshold motor activity prior to first-order responses (Gajdos, Fleming, Saez Garcia, Weindel, & Davranche, 2019). Using electroencephalography, we recently showed that a signature of motor preparation prior to first-order response correlates with confidence over different perceptual tasks (Faivre et al., 2017) and that metacognitive performance for decisions that are committed with a key press is better than that to equivalent decisions that are observed (Pereira et al., 2018). A drift-diffusion model explained this effect by showing that the accumulation of perceptual evidence is constrained by first-order decisions. Finally, transcranial magnetic stimulation (TMS) directed at the premotor cortex involved in the first-order response were found to affect confidence ratings, suggesting a causal role of action-related signals for confidence (Fleming et al., 2015).

Here, we sought to compare confidence judgements and metacognitive performance between conditions that differed only on the sensorimotor information available for the decision. To measure metacognitive performance, we designed a paradigm in which participants saw a striped visual stimulus that moved alternatively right- or leftwards for 5 s, and reported a confidence estimate regarding their capacity to discriminate the motion direction that was presented for the longest duration. Following a pre-registered plan (https://osf.io/hnvsb/), we manipulated two sources of motor information that, we hypothesized, could inform confidence judgements. First we asked participants to either observe the stimulus passively, or to track the stimulus motion direction by continuously pressing a key. Second, we asked them to report a confidence estimate after either overtly or covertly deciding about the stimulus presented for a longer time.

As per the pre-registration, we hypothesized that response-related sensorimotor activity carries information useful for confidence judgement, above and beyond the strength of the (perceptual) internal signal. We therefore expected that conditions with overt first-order responses would be associated with better metacognitive performance than those without motor responses. In the same way, we expected that conditions in which a motor action was paired with the stimulus would also be associated with better metacognitive performance than those without motor responses.

## Methods

### Participants

Twenty-seven participants took part in this study, of which four had to be excluded (see below). The results we report here correspond to a sample of 23 participants (13 males, 10 females) with a mean age (±SD) of 26.7(5). All participants had normal or corrected-to-normal vision, no colour blindness and were right-handed. Ten participants were tested in Berlin, the rest in Geneva. All received monetary compensation for their time. The procedures were approved by the corresponding local ethics committees (Ethics commission of the Institute for Psychology, HU Berlin, 2017-17 and Institutional review board of Geneva University), in conformity with the Declaration of Helsinki. Written and signed informed consents were obtained from all participants.

### Procedure

The experimental task was written in Matlab (the Mathworks, MA) using Psychtoolbox (Brainard, 1997; Kleiner et al., 2007; Pelli, 1997). Stimuli consisted of maximum-luminance red or green square (8° height and width), sinusoidal gratings (0.27 cycles/°) presented at fixation and drifting sideways at 15°/s. The green and red stimuli always drifted left- and rightwards respectively.

Each 5-second-long trial was divided into four intervals of different durations, during which four red and green stimuli were presented in alternation. The total, summed duration of each pair of same-coloured stimulus presentations corresponded to half the trial length (2.5 s) plus or minus a temporal difference determined by a staircase (see below). Further, each single stimulus presentation interval corresponded to half of the sum of the stimulus pair length.

To evaluate the effects of overt movement on metacognitive judgements, we asked for two kinds of reports: in *continuous-report* (CR+) trials, participants pressed two arrow keys using two fingers of their right hand to indicate which of the two sinusoidal stimuli was presented on the screen. In this condition, the task was simply to press a key that ‘tracked’ the motion direction of the stimulus. In conditions without continuous report (CR-), participants did not press any keys during stimulus presentation. In *trials with first-order response* (R+) trials, participants did a temporal-summation task. Upon stimulus offset, they indicated with a single key press which of the two motion directions had been presented for a longer period of time (i.e., which of the summed stimulus durations was the longest over the course of the entire 5-second trial). The response keys and hands used for the first-order response were the same as for the continuous report. In conditions without first-order response (R-), participants were also required to make a temporal summation decision (the decision was overt in R+ trials but covert in R-trials). In a factorial design, each trial corresponded to one of four possible conditions, combining continuous report (CR, “+”: present / “-”: absent) and first-order responses (R, “+”: present / “-” : absent). At the end of each trial participants rated their confidence in their decision by moving a slider with two keys on a vertical visual-analogue scale with the ends marked as “Very sure” and “Very unsure”.

The duration difference was determined separately for CR+ and CR-trials using two independent 1-up, 2-down staircases (updated only following R+ trials). We also ran two pre-experiment staircases of 25 trials each, without confidence ratings, to adjust the difference in duration of the two stimuli for each participant. After the staircases, each participant completed 240 trials in total (60 trials per condition). Trials were self-paced and the experiment took on average 50 minutes.

### Termination rule

Our plan at pre-registration was to collect data until we reach a Bayes Factor *BF_10_* of either 1/3 or 3. We started by collecting a sample of 27 participants (four excluded) and examined the data once. With this sample size we found evidence for the null hypothesis in our main test of interest (the interaction term between confidence and first-order response in the effect on accuracy as modelled by a logistic regression, see *Confirmatory analyses* below) so we halted data collection.

### Analyses

We adhered to the exclusion criteria that were pre-registered. Four participants were excluded because they did not follow the task instructions (in all cases, they did not press any keys during any of the trials in the continuous report conditions). No further participants were excluded, as none of them had first-order accuracy under 60% or above 80% in any task; and visual inspection of the staircases revealed no obvious problems. A total of 64 trials (from 17 participants) were excluded because first-order reaction times (RTs) were under 200 ms or above 5 s.

### Metacognitive performance

As per the pre-registration, we computed metacognitive efficiency (meta-d’/d’) to quantify the capacity to adjust confidence irrespective of the first-order task difficulty (Maniscalco & Lau, 2012) using the HMeta-d’ toolbox (Fleming, 2017). For that, we scaled confidence judgements for each participant by subtracting from each rating the individual minimum rating and dividing them by the total range. This procedure effectively “stretched” confidence distributions to fit the interval between 0 and 1 for all participants, thereby eliminating biases between individuals while preserving mean differences between conditions. We then discretized scaled confidence values into four confidence bins. In separate analyses, we estimated the slope parameter in a mixed effects logistic regression with accuracy as the dependent variable and confidence as the independent variable. Because mixed effects logistic regression analyses are not affected by subject-wise scaling of confidence (i.e., they include subject-wise random intercepts), we used raw confidence values as independent variables. For all models, we included a by-subject random slope for each of the main effects considered in the model, but not for their interactions. We ran Bayesian sampling of mixed regressions using the *brms* package (Bürkner, 2017, 2018) for all models, we report the estimate and its associated error M(±E) and the 95% credibility interval CI.

As no first-order responses were provided in R-trials, we defined a proxy based on the percept associated with longer key-presses during continuous report (i.e., covert first-order response). This allowed us to relate a proxy for first-order responses and confidence ratings to compute metacognitive efficiency in CR+ trials.

### Simulations for power estimations

We aimed at computing the power of our experimental design and analysis strategy. To do that, we estimated the proportion of simulated “experiments” in which we would have found a significant difference between two given conditions with different M-ratios. We used signal detection theory to simulate first and second-order responses from 80 trials for each of the 23 participants (see Figure 4, A). We set the distributions of the internal signals elicited by the stimuli to be a normal distribution with μ = ±d’/2, σ = 1 (the sign of μ depended on the longer stimulus presented). First-order responses were defined according to an optimal first-order criterion at 0.

**Figure 1:**
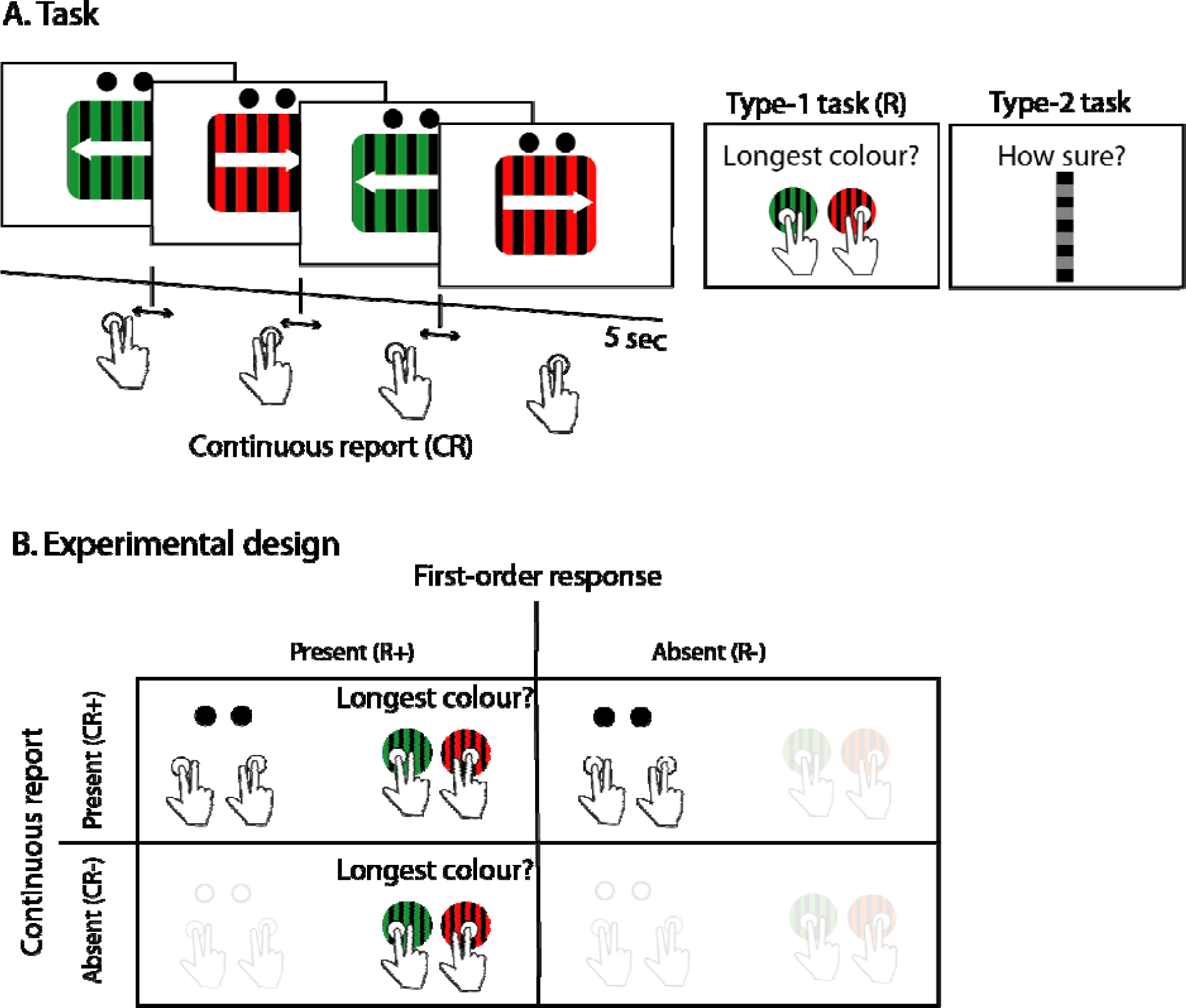
**(A.) Task.** Example trial with both continuous report (CR+) and first-order response (see panel B.) On each 5-second trial, two stimuli pairs appeared serially in four consecutive intervals. Participants pressed one of two keys for the entire duration of the trial, tracking the visual presentation (continuous report). Following stimulus offset, participants reported which of the two stimuli had the longest duration overall. **(B.) Experimental design.** Trials followed a 2×2 factorial design that combined continuous report and first-order responses (trials could therefore be one of the four possible conditions, CR+R+; CR+R-; CR-R+; or CR-R-). Participants rated their confidence in all conditions. Thus, the task demanded that participants make a first-order judgement in every trial, but the corresponding overt action was only present in R+ conditions.

**Figure 2:**
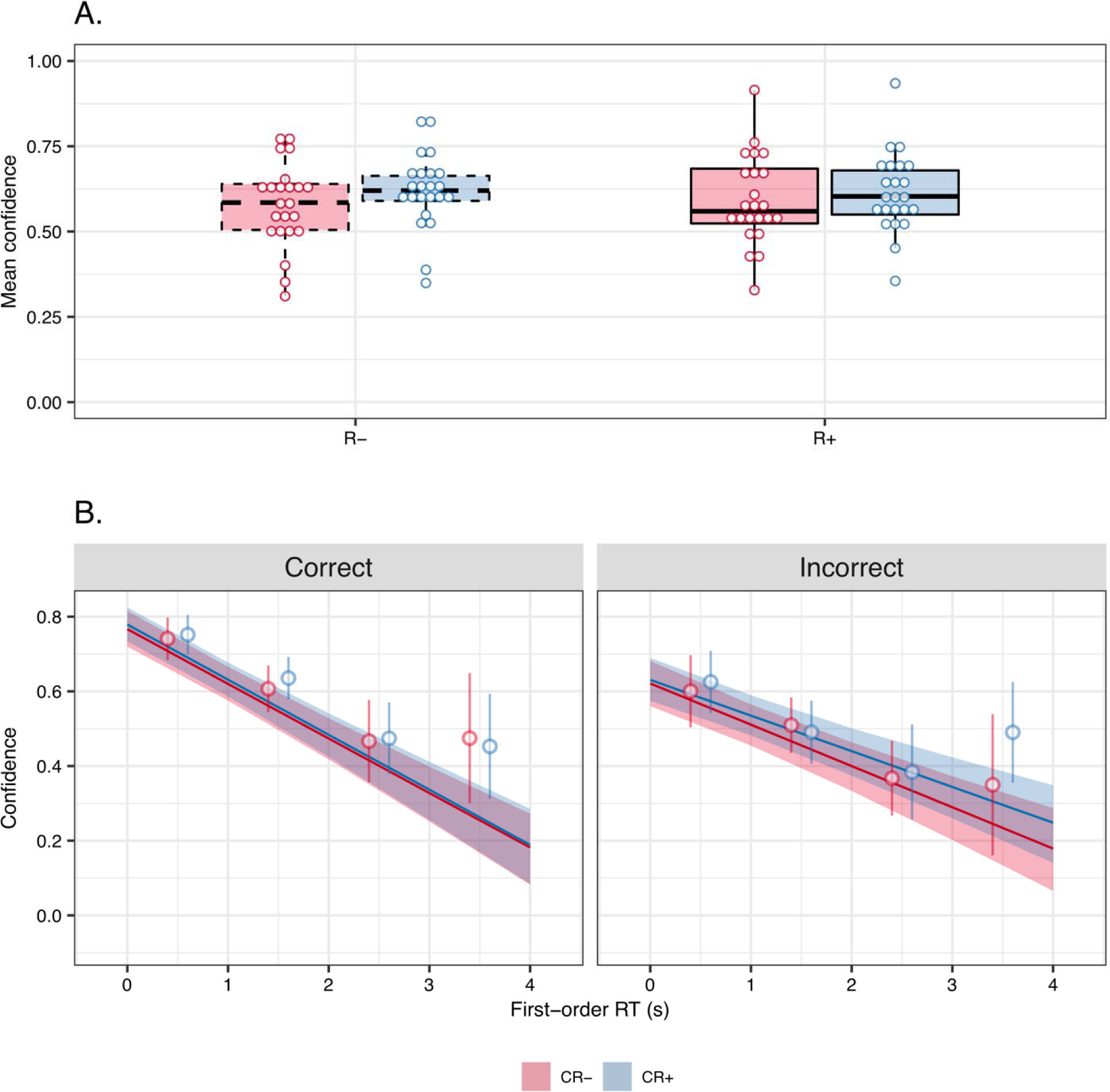
**(A.) Differences in confidence judgements between conditions.** Trials with continuous report (CR+) were associated with higher confidence. **(B.) Figure 2: relationship between first-order reaction times and confidence judgements.** As expected, confidence judgements had a strong negative relationship with first-order reaction times. This relationship was present in all R+ trials (R-trials were not included in this analysis) but was stronger in the subset of correct trials. Regression lines and confidence intervals around them represent the model fit. The model took continuous reaction times as input. For illustrative purposes, we plot open circles and error bars that represent mean ± 95%CI over participants after rounding reaction times and subtracting 0.5 s.

**Figure 3:**
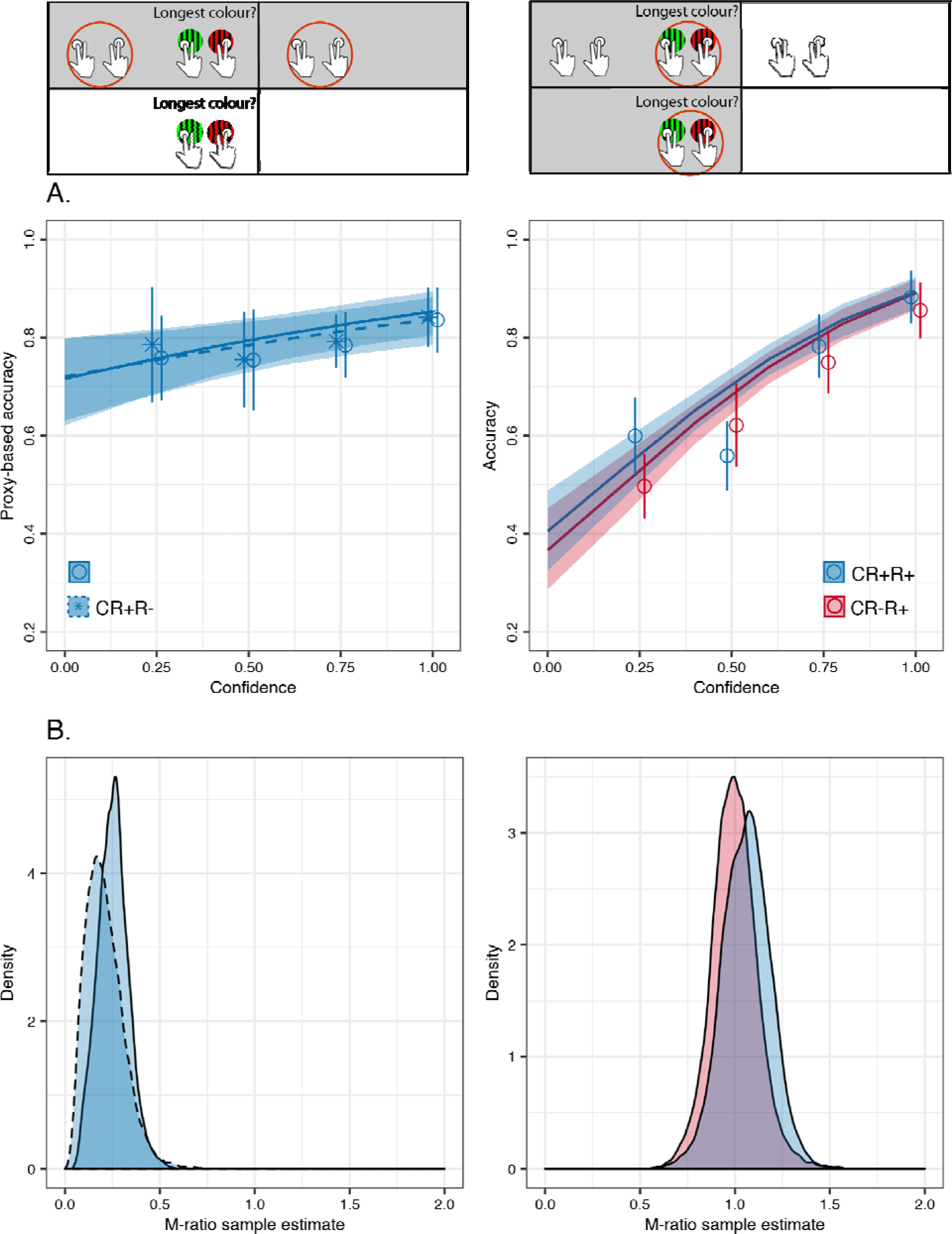
Differences in metacognitive performance between conditions. **(A.) Metacognitive sensitivity quantified with a regression model on accuracy vs. confidence.** Estimated regression curves from the proxy for first-order response (left panel) and overt first-order response (right panel). The presence of a first-order response did not affect the relationship between confidence and the proxy’s first-order accuracy. Open circles and error bars represent mean ± 95%CI over participants after rounding confidence ratings. **(B.) Metacognitive efficiency quantified with M-ratio.** As in panel A., we found no evidence that giving either an overt first-order response (left panel) or that pairing an action to perceptual input (right panel) improved metacognitive efficiency. The insets above the panels highlight (in grey) which trials were used for each of the analyses.

**Figure 4:**
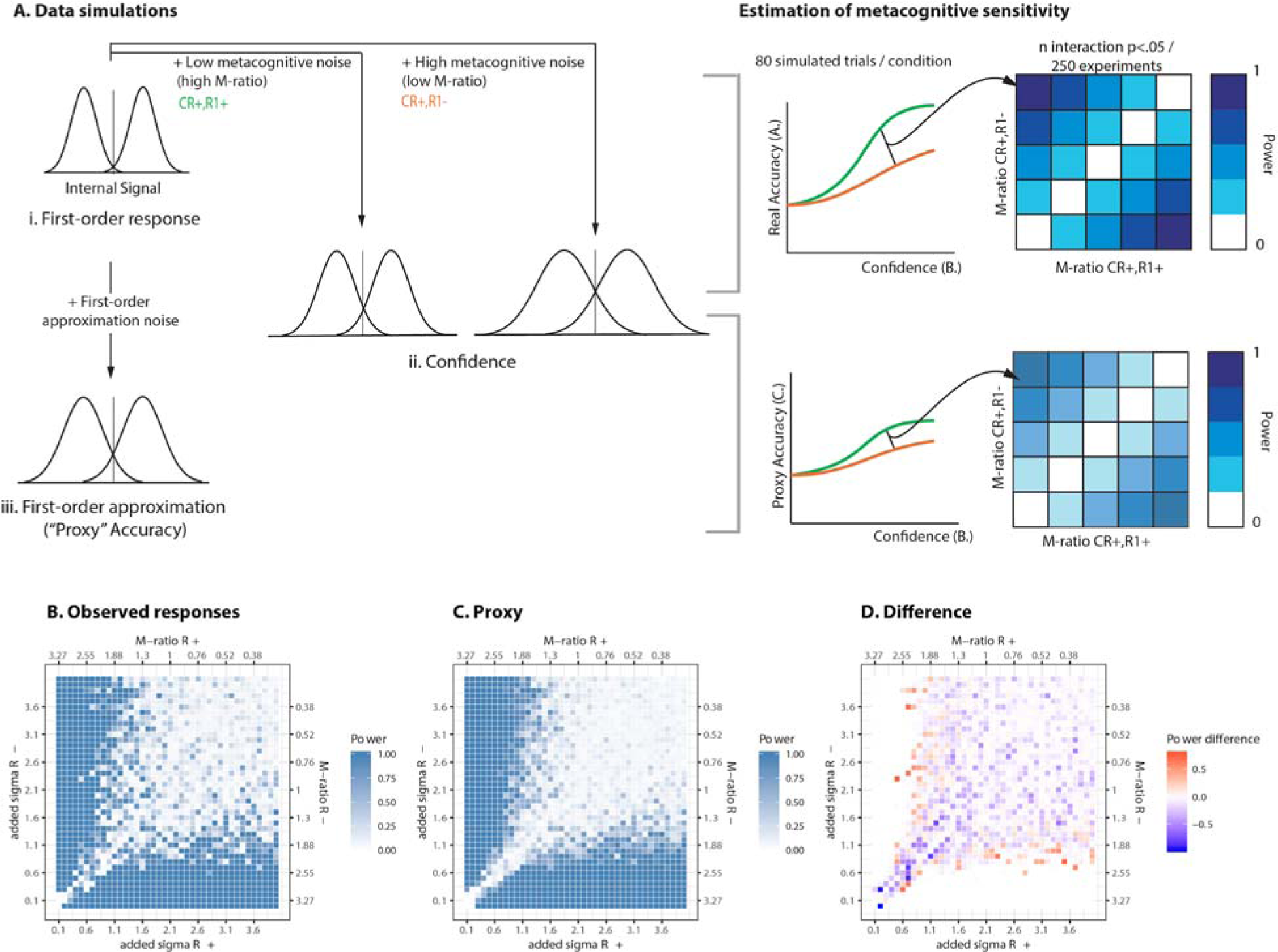
Power simulations. **(A.) Data simulation strategy.** We considered two conditions (in this case, CR+R+ and CR+R-), expected to differ in M-ratio. For each one of 250 experiments, we simulated 80 trials per condition, drawing three values for each of the “real” internal signal (top left, *i.*), a noisy confidence estimate (internal signal + metacognitive noise for each of two conditions, middle row *ii.*) and a value for the noisy proxy (bottom left, *iii.*). We fed the simulated trials into a logistic regression model, and determined the power of our analysis, i.e. the proportion of “experiments” in which the interaction term (representing a difference in metacognitive sensitivity between conditions) was significant (right). **Results.** Power estimations for the analysis based on actual responses **(B.)** and for proxy-based responses **(C.).** Panel **(D.)** shows the power difference between (B.) and (C.). There are no differences in power when differences in M-ratios between two conditions are large (regions away from the diagonal) whereas there are small decreases in power for the proxy-based analysis for combinations of M-ratios that are closer to the diagonal.

Next, to simulate the first-order proxy, we injected randomly distributed noise into the internal signal, sampled from a normal distribution centered at μ = 0 and σ = 0.8. This led to a correspondence of approximately 70% between real and proxy simulated responses, similar to our data. The rationale for adding noise to the internal signal rather than to the binary response variable itself was to preserve the structure of the data: trials with an internal signal closer to the decision boundary are associated with lower confidence, and therefore are more likely to cross-over the decision boundary as a consequence of adding noise, compared to trials with an internal signal strength that is far from the decision boundary. We obtained the simulated proxy by binarizing the noisy internal signal data based on the position relative to the same optimal first-order criterion placed at 0.

Finally, to simulate confidence ratings, we first added noise to the simulated internal signal, sampled from a normal distribution centered on 0 and with σ ranging between 0 and 4. To simulate M-ratio values above 1, we then swapped the identity of the two distributions, in order to make the second-order distributions sharper than the first-order ones. In a separate simulation, we established that these values of added σ corresponded to M-ratio values ranging between 0 and 1.1, which corresponds to the range of M-ratios in our experimental data (see figure 3). We set the simulated confidence as the absolute value of the internal signal, that is the distance to the first-order decision criterion. Thus, we added two kinds of noise to the original internal signals, with different meaning: The first type of noise simulated the imperfect relationship between covert/overt responses and their corresponding proxy. The second type of noise simulated the imperfect mapping between the strength of the internal signal at the point of the first- and second-order decisions, a relationship captured by M-ratio (Maniscalco & Lau, 2012).

We then submitted these simulated data (for 80 trials from 23 participants) to the same mixed effects logistic regression we used to analyze empirical data. We repeated this procedure 250 times with each combination of M-ratio, to estimate the number of times that a significant effect would occur in 250 experiments.

## Results

### Descriptive analyses – effects on confidence

The adaptive staircase procedures successfully fixed performance at approximately 71% correct: mean accuracy (±SD) was 72.0(4.2)% for continuous report and 72.1(4.6)% for no continuous report conditions, with no difference between conditions (*t*(22) = −0.12, p = 0.91, d = −0.02, *BF_10_* = 0.22). Mean perceptual evidence did not differ across CR+R+ and CR-R+ trials (t(22) = 1.75, p = 0.09, d = 0.36, *BF_10_* = 0.54), indicating that pairing motor information to the perceptual input was not informative for the first-order decision. Next, we tested for mean differences in confidence between all conditions using a linear mixed effects regression model on confidence. The model included the two experimental manipulations (R and CR) and their interaction as fixed effects, intercepts for subjects as random effects, and a by-subject random slope for each of the factors. We found no interaction between overt first-order and continuous report (M = −0.02(0.01), evidence ratio = 0.10), no strong effect of overt first-order responses on mean confidence ratings (M = 0.01 (0.02), evidence ratio = 3.43), but a significant increase of mean confidence in conditions with continuous report (M = 0.04(0.02), evidence ratio = 75.92) (Figure 2A).

Importantly, to test the hypothesis that the monitoring of first-order responses or their underlying processes contributed to confidence, we first established the existence of a relationship between reported confidence and first-order RT. We did so by fitting a mixed effects linear regression to confidence in trials with overt first-order responses (R+), including first-order accuracy, first-order RT, condition (CR+/CR-), and perceptual evidence as fixed effects, random intercepts for subjects, and by-subject random slopes for each fixed effect. As expected, we found a strong main effect of first-order RT on confidence (M = −0.15(0.02), evidence ratio > 4000), confirming the relationship that has been reported in previous studies (e.g. (Fleming et al., 2010; Patel et al., 2012) (Figure 2B). This effect was stronger for correct trials than for incorrect trials (interaction effect estimate M = 0.04(0.02), evidence ratio = 46.06). We also found a main effect of accuracy (M = −0.15(−0.03), evidence ratio > 4000) and of perceptual evidence (M = 0.22(0.06), evidence ratio > 4000), indicating that confidence was higher for correct responses, and in the presence of higher perceptual evidence. However, the model revealed no main effect of condition (M = 0.01(0.02), evidence ratio = 0.41). No other model parameters were associated with confidence.

Together, these results indicate that fast first-order responses were associated with higher confidence, but that response times are unlikely to play a causal role as removing first-order responses altogether had no effect on confidence.

### Confirmatory analyses – effects on metacognitive sensitivity

Our first hypothesis was that sensorimotor activity related to first-order responses carries information useful for confidence, above and beyond the strength of perceptual evidence. We therefore expected conditions with overt first-order responses to be associated with better metacognitive sensitivity (measured as the relationship between confidence and first-order accuracy) than those without motor responses. As we could not calculate first-order response accuracy in conditions with no first-order responses (R-), we assumed that the percept associated with longer key-presses during continuous report corresponded to the covert first-order response. In CR+R+ trials, this proxy based on continuous report predicted the actual first-order response in 65.5(8)% of trials (ranging between 50% and 79.6%). For CR+R+ trials, we confirmed that response predictability based on the stimulus (i.e., longest stimulus presented) and proxy (i.e., key pressed the longest) was significantly higher than based on the stimulus alone (difference in Bayesian information criterion = 2.9; □^2^ = 10.04, p = 0.002). This is why despite low predictability scores, we proceeded with this analysis as per our pre-registered plan, and pursued alternative ways to analyze the data in Exploratory analyses below.

To compare metacognitive sensitivity between conditions with and without first-order responses, we built a mixed effects logistic regression for proxy accuracy that included condition (CR+R+/CR+R-) and confidence and their interaction as fixed effects, as well as subject-wise random intercepts, and random slopes for both confidence and condition. If metacognitive monitoring is affected by the presence of first-order responses, this should manifest as a significant interaction effect between confidence and the presence of a first-order response: the relationship (slope) between confidence and proxy accuracy should be stronger for trials with first-order responses than for those without them. Against our expectations, but in line with the results on mean confidence reported above, we found no interaction effect (M = −0.11(0.39), evidence ratio = 1.57). On the other hand, a main positive effect of confidence (M = 0.82(0.32), evidence ratio = 116.65), indicated that the likelihood that the proxy was the correct answer increased with confidence and thus, simply put, that participants had some metacognitive access to their response accuracy. The estimation of M-ratio (meta-d’/d’) revealed consistent results, as we found no differences between conditions in the M-ratio estimates (R+: M-Ratio = 0.22, HDI = [0.12 0.42], R- : M-Ratio = 0.25, HDI = [0.10 0.45]; difference between conditions: HDI = [-1.42 0.89]).

Our second pre-registered hypothesis was that metacognitive performance between conditions with and without continuous report would differ, because the key presses in the continuous report constitute an additional source of information for confidence responses. To test this hypothesis, we followed two approaches. First, using the same approach as above, we measured metacognitive sensitivity as the relationship between confidence and first-order accuracy. Here again, a main effect of confidence on accuracy (M = 2.51(0.37), evidence ratio > 4000) suggested that participants could monitor their performance. However, we found no interaction between confidence and condition (M = 0.13(0.35), evidence ratio = 1.77), indicating that this effect was comparable with and without continuous report. This analysis included only trials with overt first-order responses, so it was possible to measure metacognitive accuracy with standard methods. Thus, we also estimated M-ratio (meta-d’/d’) in trials with and without continuous report. Again, and consistent with our regression analyses, we found no differences between conditions in the M-ratio estimates (CR+ : M-Ratio = 1.06, HDI= [0.83 1.32], CR- : M-Ratio = 0.98, HDI = [1.24 0.77]; difference between conditions: HDI = [-0.27 0.40]).

Our data revealed no differences in the relationship between confidence and first-order accuracy between conditions.

To measure the effect of first-order responses (CR+R+ vs CR+R-) we relied on a proxy as the best informed guess for the covert first-order response; but the proxy was noisy and corresponded to the overt first-order response only for approximately 65% trials over all participants. In other words, with this analysis we injected noise into our first-order response, which might have in turn affected both the value of the confidence × condition interaction estimates and our ability to find robust effects. To examine whether this was the case, and to what extent this affected our results, we simulated data from 250 “experiments” to compare the power of the logistic regression analysis based on the simulated first-order response and on the degraded first-order proxy.

The results of these simulations (Figure 4.B-D) validated our analysis strategy. First, we found that power between the two analyses did not differ for values far from the diagonal (i.e., pairs of M-ratios with large differences between them). Second, and crucially, we found that even in regions where the proxy analysis fared worse (i.e., had lower power), power reductions were only marginal, in the range of 0.1-0.3, and are thus unlikely to affect our results fundamentally. Interestingly, power estimations for the proxy-based analyses showed a somewhat smoother pattern than those from actual responses. This result, presumably an effect of having an additional source of Gaussian noise, may be an unexpected advantage of the proxy-based analysis in preventing false inferences.

### Exploratory analysis – machine learning tools to predict first-order responses

Finally, we considered that the relatively low predictability of the continuous report-based proxy could be poor due to its simplicity: the proxy was based on nothing more than the longest reported percept in each CR+ trial. To extract as much information as possible from CR+ trials, we leveraged standard machine learning (ML) algorithms to predict first-order responses from CR+ information. First, for each CR+R+ trial we extracted features including the number of transitions in the key press response, the identity of the first and last stimuli shown and keys pressed, the total time with correct and incorrect key presses, and the delay between each stimulus presentation and the response. Using the scikit-learn module in Python (Pedregosa et al., 2011), we then trained three different classifiers on the data pooled over all participants using leave-one-out cross-validations: logistic regression, naive Bayes, and k-nearest neighbours. Their accuracy, based on the confusion matrix on CR+R+ trials revealed low overall predictability: 0.63, 0.61 and 0.64 respectively. These relatively low values are comparable to those of our simple proxy and we therefore did not carry out any further analyses with the ML-based predictions.

## Discussion

The past years have seen a growing interest in elucidating the sources of information that contribute to confidence judgments, as a window into potential computational processes that allow the brain to monitor itself. Converging evidence from very different experimental paradigms suggested that confidence is modulated by motor information concurrent with the first-order response (Faivre et al., 2017; Fleming et al., 2015; Patel et al., 2012; Pereira et al., 2018); for a review see (Anzulewicz, Hobot, Siedlecka, & Wierzchoń, 2019). Here, we set out to directly investigate this possibility. Concretely, we used a temporal-summation metacognitive task and asked whether committing a motor response associated with the response affected corresponding confidence judgements.

### Effect of first-order responses on confidence ratings

As a precondition for our analyses, we first replicated what several studies had shown before (e.g., (Fleming et al., 2010; Patel et al., 2012): in trials with overt first-order responses (R+), reaction times to the first-order task showed a clear negative correlation with reported confidence. Based on these results alone, our data are in principle compatible with the hypothesis that first-order responses influence reported confidence. Crucially, we tested this hypothesis directly by comparing two conditions of the task that differed in whether participants had overtly responded with a key press to the first-order task (CR+R+), or if their response remained covert (CR+R-). We first compared conditions in terms of average confidence judgements. Against our expectations, and despite the strong correlation between first-order reaction times and confidence, we found that absolute confidence judgements did not vary with the presence or absence of overt responses.

To further investigate the effects of overt responses, we then examined an important aspect of confidence judgements, namely their precision. That is, we considered that while participants may not have felt in general less confident in trials with covert responses, the quality of confidence judgements might have been degraded, resulting in a decrease in metacognitive performance relative to trials with overt responses. Measures of metacognition (metacognitive *sensitivity*, based on logistic regression and *efficiency*, based on M-ratio) rely on relating trial-wise confidence to accuracy. As the identity of covert responses remained latent by design, we inferred them relying on a proxy based on continuous reports (CR+). Concretely, we considered the percept with the longest key press as a proxy for both overt and covert first-order responses. We then compared metacognitive sensitivity and efficiency based on the relationship between confidence and the proxy for responses. Here, mirroring the results from the analysis of absolute confidence values, we found no effect of overt first-order responses. A concern with this analysis is that the proxy only corresponded to actual (overt) responses in an average of approximately 65% of trials, which resulted in a systematic under-estimation of metacognitive performance (e.g., compare dashed blue lines between panels in Figure 3). However, as the proxy predictive power does not differ across conditions, comparisons of metacognitive performance across conditions are still legitimate. To further assess the effect of this poor predictability on our conclusions, we ran power estimations based on simulated data. These simulations successfully reproduced the observed effects, and revealed that power reductions were only minimal.

### Effect of continuous report on confidence ratings

In our factorial design, we also tested for the effect of continuous report paired to stimulus presentation on confidence judgements. Over conditions with and without first-order responses (both R+ and R-), we found a consistent increase in confidence following continuous report (CR+ vs. CR-). Under one simple account, this effect could be explained by attentional demands: The requirement to pair voluntary key presses to the stimuli could have led participants to attend to the visual stimuli more strongly in CR+ trials as compared to CR-trials. While we cannot rule out this explanation, we note that attentional differences should have also led to a perceptual advantage. Against this prediction, we found no differences in the values of duration difference (i.e., task difficulty) resulting from the staircase. Thus, there is no clear evidence that simple differences in attentional demands between conditions could account for participants’ higher confidence ratings. Alternatively, higher confidence ratings may result from criterion shifts. In fact, our model comparison showed that motor behaviour could explain first-order choices over and above perceptual evidence, suggesting that key presses in continuous report conditions were an additional source of information available for both the perceptual (first-order) and the confidence (second-order) tasks. With additional sources of information, participants may place their second-order criteria more liberally, resulting in higher confidence ratings.

### Differences with the existing literature

To the best of our knowledge, this study is unique in that the effect of motor components on confidence was investigated by completely *removing* the first-order response in some conditions, and replacing it instead with actions paired to the stimuli presentation. As a consequence, we never required participants to provide explicit responses in covert-response conditions. Instead, we inferred them through participants’ continuous report. Other studies have addressed the same question by using different experimental manipulations, that can be broadly grouped as following one of three approaches. A first set of studies have asked participants to rate confidence of *observed*, rather than committed, actions, whilst letting participants observe only first-order RTs (Patel et al., 2012; Vuillaume, Martin, Sackur, & Cleeremans, 2019) or both RTs and stimuli (Pereira et al., 2018) before making the confidence judgement. A second group of studies have instead manipulated the timing of the confidence judgement relative to that of the first-order response (Siedlecka, Paulewicz, & Wierzchoń, 2016; Wokke, Achoui, & Cleeremans, 2019). Finally, a third approach consists of directly manipulating motor signalling either physiologically using TMS (Fleming et al., 2015) or behaviourally by instructions (Faivre et al., 2018; Palser, Fotopoulou, & Kilner, 2018). Here, we followed the novel strategy of removing first-order responses and instead inferring them from stimulus-coupled responses. Against what has been reported in the literature and our expectations, we found that bypassing first-order responses had no observable effect on metacognitive performance.

Our results also revealed that continuous motor responses contingent to perceptual evidence significantly increased confidence. A brief review of the literature reveals that motor activity impacts confidence biases and metacognitive performance distinctively, with large variations across experimental paradigms. On the one hand, our results are in line with what was reported by Gajdos and colleagues (2019), who found that subthreshold motor activity prior to a decision increased confidence bias, with no impact of metacognitive performance. Other experimental manipulations produced the converse effect, namely a modulation of metacognitive performance with no change in confidence bias. This includes the comparison of confidence in committed vs. observed decisions (Pereira et al., 2018), and confidence under high or low sensorimotor conflicts (Faivre et al., 2018). Using a similar design comparing prospective and retrospective confidence judgments, Siedlecka and colleagues (Siedlecka et al., 2016) found that both confidence bias and metacognitive performance increased in presence of action-related signals. This set of mixed results questions the functional relevance of motor signals, and suggests that the relationship might be more complex than previously thought. We speculate that the computation of confidence may be flexible, and largely depend on the information that is globally available. In all previous studies, to the best of our knowledge, participants had access to some form of first-order reaction time information, at some point in time during the trial: either through observation from the third-person perspective, directly after the confidence report or through simple access to reaction times produced under experimentally manipulated motor signals. In our no-report conditions, instead, responses were completely absent and may have shifted participant’s global strategies for the computation of confidence. In other words, we contest that while first-order reaction time information is, under some experimental settings used by participants to generate a confidence judgement, when motor information is not available at all, it may be replaced by other, equally precise sources of information, closer to the strength of evidence (such as the probability of being correct (Sanders et al., 2016), the internal signal noise (Navajas et al., 2017) and the evidence in favour of the chosen response alternative (Peters et al., 2017). This admittedly speculative account is compatible with our capacity to form confidence estimates about decisions that are not directly linked to a transient motor action, for instance when controlling a brain machine interface (Schurger, Gale, Gozel, & Blanke, 2017) or when making global confidence judgments in ecological contexts (Rouault, Dayan, & Fleming, 2019).

### Limitations and future directions

A limitation of our design lies in the capacity to identify covert first-order responses from continuous reports. While voluntary key presses paired to the stimuli shown on the screen were a relatively poor predictor of covert responses, we argue that the approach is promising given that future lines of research might take this first step further in order to develop ‘no-report’ paradigms where covert decisions can be unequivocally inferred without a margin for error. Potential approaches include either eliciting an automatic response like the optokinetic nystagmus (Frässle, Sommer, Jansen, Naber, & Einhäuser, 2014), instead of a voluntary one like the key presses we used here; requiring voluntary key presses in highly trained participants, leading to low latencies between perception and response; or inferring responses through covert attention measured using steady-state visual evoked potentials (SSVEP, e.g. (Heering, Beauny, Vuillaume, Salvesen, & Cleeremans, 2019)). Another limitation is the use of adaptive staircase procedures throughout the experiment. While maintaining task difficulty constant across trials, conditions, and participants is important to finely estimate metacognitive performance (Rahnev & Fleming, 2019), it also may hinder the relevance of sensorimotor signals as informative cues regarding the difficulty with which a decision was made (Kiani, Corthell, & Shadlen, 2014). Thus, a possibility is that sensorimotor signals are more potent cues for confidence estimates under fluctuating task difficulty. Notwithstanding these limitations, we note that we found a clear null effect on absolute confidence differences between conditions with covert and overt first-order responses. This result, which is not contaminated by imprecision in our identification of covert first-order responses, more strongly argues for our interpretation that motor signals need not be used in metacognitive monitoring.

### Conclusion

Identifying the sources of information that feed into confidence judgements is a core issue in metacognition research. This study suggests that, while confidence judgements correlate with first-order reaction times, this relationship may be merely correlational, as removing the execution of first-order decisions altogether had no visible impact on confidence nor metacognitive performance. By contrast, motor actions paired to stimulus presentation boosted confidence, but not metacognitive performance. These results, then, do not support the emerging idea that metacognition relies on the monitoring of sensorimotor signals, and call for further research to find the underpinnings of metacognitive judgements.

## Acknowledgments

We thank Lukas Röd, Carina Forster, and Marco Wirthlin for help with data collection. We also thank Guillermo Bernabó for help with machine learning analyses, and Michael Pereira and Matthias Guggenmos for helpful comments on an earlier version of this manuscript. EF and CK are supported by a Freigeist Fellowship to EF from the Volkswagen Foundation (grant number 91620) and by the Deutsche Forschungsgemeinschaft (DFG, German Research Foundation) – 337619223 / RTG2386. NF is supported by a Starting Grant from the European Research Council (#803122).

## Contributions

EF and NF developed the study concept and contributed to the study design. EF and CK wrote the experimental scripts. Data collection was performed by CK. Data analysis was performed by EF, CK and NF. EF and NF drafted the paper; all authors provided critical revisions and approved the final version of the paper for submission.

## Links

The pre-registered analysis plan can be found at: https://osf.io/hnvsb/. The raw data, analysis files to reproduce all figures are available under https://gitlab.com/nfaivre/filevich_metareport.

